# Towards CNN Representations for Small Mass Spectrometry Data Classification: From Transfer Learning to Cumulative Learning

**DOI:** 10.1101/2020.03.24.005975

**Authors:** Khawla Seddiki, Philippe Saudemont, Frédéric Precioso, Nina Ogrinc, Maxence Wisztorski, Michel Salzet, Isabelle Fournier, Arnaud Droit

**Affiliations:** Centre de Recherche du CHU de Québec - Université Laval, Québec City, QC, Canada; Univ. Lille, Inserm, CHU Lille, U1192 - Protéomique Réponse Inflammatoire Spectrométrie de Masse - PRISM, F-59000 Lille, France; Université Cote d’Azur, CNRS, INRIA, I3S, Sophia Antipolis, France

**Keywords:** CNNs, Transfer Learning, Cumulative Learning, Classification, Small Datasets

## Abstract

Rapid and accurate clinical diagnosis of pathological conditions remains highly challenging. A very important component of diagnosis tool development is the design of effective classification models with Mass spectrometry (MS) data. Some popular Machine Learning (ML) approaches have been investigated for this purpose but these ML models require time-consuming preprocessing steps such as baseline correction, denoising, and spectrum alignment to remove non-sample-related data artifacts. They also depend on the tedious extraction of handcrafted features, making them unsuitable for rapid analysis. Convolutional Neural Networks (CNNs) have been found to perform well under such circumstances since they can learn efficient representations from raw data without the need for costly preprocessing. However, their effectiveness drastically decreases when the number of available training samples is small, which is a common situation in medical applications. Transfer learning strategies extend an accurate representation model learnt usually on a large dataset containing many categories, to a smaller dataset with far fewer categories. In this study, we first investigate transfer learning on a 1D-CNN we have designed to classify MS data, then we develop a new representation learning method when transfer learning is not powerful enough, as in cases of low-resolution or data heterogeneity. What we propose is to train the same model through several classification tasks over various small datasets in order to accumulate generic knowledge of what MS data are, in the resulting representation. By using rat brain data as the initial training dataset, a representation learning approach can have a classification accuracy exceeding 98% for canine sarcoma cancer cells, human ovarian cancer serums, and pathogenic microorganism biotypes in 1D clinical datasets. We show for the first time the use of cumulative representation learning using datasets generated in different biological contexts, on different organisms, in different mass ranges, with different MS ionization sources, and acquired by different instruments at different resolutions. Our approach thus proposes a promising strategy for improving MS data classification accuracy when only small numbers of samples are available as a prospective cohort. The principles demonstrated in this work could even be beneficial to other domains (astronomy, archaeology…) where training samples are scarce.

## 1 Introduction

Accurate and rapid identification of cancer tissues has a crucial impact on medical decisions. Conventional histopathological examinations are resource intensive and time-consuming, requiring 30–45 minutes per sample processed and the presence of a skilled pathologist [1]. A similar need exists in the treatment of infections, where accurate identification of microorganisms responsible for human infection is important to ensure the most appropriate and effective treatment for a patient, in the shortest possible time [2]. In this context, it is essential to use methods which provide accurate identification and correct interpretation of the analyzed samples. Mass spectrometry (MS) is particularly useful for such purposes since it provides non-targeted molecular information on the millisecond time scales. Its sensitivity, reproducibility, and suitability for analyzing complex mixtures are well established. New analysis methods of crude samples are making diagnosis even faster and easier. Simultaneously, the development of MS-based bacterial biotyping clearly illustrates the value of MS in rapid clinical applications [3].

For cancer-related diagnosis and microbial pathogen identifications, many popular classification Machine Learning (ML) models, such as Support Vector Machine (SVM) [4], Random Forest (RF) [5], and Linear Discriminant Analysis (LDA) [6] have been already used and compared [7–10]. These ML methods are applied to preprocessed MS data, and differences in preprocessing pose a major challenge to any comparison of MS data analysis. Classification model design for rapid applications thus becomes a highly complex task, since it must follow a workflow involving several interdependent preprocessing steps. Data preprocessing is used to improve the robustness of subsequent multivariate analysis and to increase data interpretability by correcting issues associated with MS signal acquisition [11]. Preprocessing quality is important, and if inadequate, can lead to biased or biologically irrelevant conclusions [12]. Several factors, often related to the experimental conditions including sample heterogeneity, sample processing and MS analysis (e.g. electronic noise, instrument calibration stability, temperature stability,…) and other experimental conditions can contribute to spectral variations including shifts in peak location, fluctuating intensities and signal distortion [13]. In other words, peaks corresponding to the same molecule in different samples can be shifted and their signal intensity can vary from one spectrum to another [14–16]. Signals of lower intensity are in general more affected by such variations because they can become buried in baseline noise in certain cases. Since these include many markers of interest, this may lead to loss of important biological information [17]. Corrections on peak position variations are required in order to align different spectra properly and thus ensure consistency in downstream analysis. This alignment constitutes a significant hindrance to achieving reproducibility especially in today’s complex datasets, and remains a challenging problem since it is neither linear nor uniform across the whole collection of MS spectra [17]. In addition to peak shifts, other spectral fluctuations must be corrected in order to minimize background and serious intensity distortion due to noise and baseline drift caused by instrument electronics, ion saturation or contaminants within the samples [13]. To overcome batch effects, peak intensities must be equalized to reduce overall signal variation between acquisitions using intensity calibration or normalization [18, 19]. Log-intensity transformation is one of the methods most commonly used to attenuate large differences in variability differences between peaks across the spectrum [19]. Another preprocessing step that is crucial to subsequent analysis is the peak detection, also known as peak picking. This consists of identifying informative peaks that correspond to a true biological signal by finding all local extremes in the spectrum, which corresponds to the conversion of spectra from profile to centroid mode [20]. Finally, the *curse of dimensionality*, must be avoided. This is a well-known problem that arises when processing MS data having a large number of dimensions, and is lessened using data dimensionality reduction techniques [21]. Various MS classification workflows have been developed so far, but there is no golden standards for the optimal choice of parameters at each individual step, for their quality evaluation or for their best combination [22]. It has been shown that the choice of preprocessing parameters for a specific dataset can decrease the performance of the classification model and that preprocessing may be effective only for that dataset and not any others generated from different instruments or with different settings [23]. A standard pipeline for MS classification using SVM, RF or LDA must include these preprocessing steps and must consider aforementioned constraints, which makes such algorithms unsuitable for rapid analysis.

Convolutional Neural Networks (CNNs) are one of the most successful deep learning architectures designed to learn representation from an input signal with different levels of abstraction [24]. A typical CNN includes convolutional layers, which learn spatially invariant features from input (i.e. invariance to translation, invariance to scale, etc) stored in feature maps, pooling operators that extract the most prominent structures, and fully connected layers for classification [25]. To address rapid clinical MS data classification tasks, CNNs represent an attractive approach offering various advantages over conventional ML algorithms. These include significantly higher accuracy, effectiveness on raw spectrum classification even in presence of signal artifacts (noise, baseline distortion, etc.) and hence discards the need for data preprocessing before classification [26], integration of features extraction with classification and without a feature-engineering step since all layers are trained together, and finally exploitation of spatially stable local correlations by enforcing the local connectivity patterns, where the output of each layer of these networks is directly related to small regions of the input spectrum [27]. However, CNNs classification efficiency trained using a small number of spectra drops rapidly [26]. Unfortunately, many real-world applications do not have access to big training sets because of data scarcity, or because of the difficulty and expense in labeling data [28]. In medicine, it is often the case that some samples are only accessible in limited amounts, especially for rarer diseases and pathologies (e.g. patient biopsies, at advanced stage of infection, etc). Therefore the size of clinical datasets is constrained by data availability and by the experiments complexity and high cost [29]. For such applications, transfer learning has emerged as an interesting approach [30]. This technique is applicable to small datasets and therefore requires fewer computational resources while increasing the classification accuracy as compared to CNNs models built from scratch. Transfer learning is a two-step process. An accurate data representation is first learned, by training a model on a dataset containing a large amount of annotated data covering many categories. This representation (i.e. its model weights) is then reused to build a new model based on a smaller annotated dataset containing fewer categories, by training only the final decision layer(s) or by also fine-tuning the whole model with the reduced set of categories. Transfer learning has proven useful in many engineering areas including computer vision, robotics, image classification and natural language processing (NLP) applications [31]. With MS data, it would use basic similarities in spectral shape gathered from different datasets and adapted to address new classification problems. This has yet to be explored for 1D spectral data, since no 1D spectral dataset as large as the ImageNet database in the 2D image analysis domain is available [32]. Most of MS classification by CNNs focused on MS 2D imaging analysis [33–35]. Only few studies of input signal classification or regression using 1D-CNNs with vibrational spectroscopy data [36], Near-Infrared (NIR) spectroscopy data [37–39] or Raman spectroscopy data [26] have been published. We have found no description of their use or of transfer learning or representation learning in conjunction with 1D-MS data. The aim of this study is to build CNNs-based classification models for 1D mass spectra by transfer learning or representation learning. Pattern recognition models are built using small clinical datasets generated for the diagnosis of cancers or microbial infections. Our work bridges the gap between developing a novel CNNs framework and its application on small datasets classification. The approach is fully applicable to other domains where the lack of data is still a hindrance.

## 2 Methods

### 2.1 Datasets

These independent MS datasets are used to evaluate the proposed approaches :

1. The **canine sarcoma dataset** contained 1 healthy and 11 sarcoma histology types obtained from the 33 annotated *ex vivo* biopsies as described previously [40]. Spectra are acquired in sensitivity positive ion mode using a Synapt G2-S Q-TOF MS instrument (Waters, Wilmslow, United Kingdom). The multi-classification model presented is focused on sarcoma type only. Tumor type grading is beyond the scope of this study.
2. The **microorganism dataset** contained a five human pathogen collection of 2 Gram-negative (Gram-) bacteria, 2 Gram-positive (Gram+) bacteria, and 1 Yeast cultivated as described previously [41]. Spectra are acquired in positive high-resolution mode using a Synapt G2-S Q-TOF MS instrument (Waters, Wilmslow, United Kingdom). SpiderMass is a new system designed for mobile *in vivo* and real-time surface analysis. The instrumentation setup is described in detail elsewhere [42]. SpiderMass does not involve a chromatographic step, making it compatible with rapid analysis but increasing output spectrum heterogeneity. These two small SpiderMass datasets are characteristic of the clinical field, where samples availability coming from patients can be limited thus making the task of classification models more difficult.
3. The MALDI-MSI rat brain dataset contained spectra of rat gray and white brain matter, acquired using a Rapiflex MALDI-TOF instrument (Bruker, Bremen). MALDI-MSI mass spectrometry (MALDI-IMS) are imported into the user-friendly Scils software (Bruker Daltonik GmbH) and ROI non-processed spectra are exported into a csv file format.
4. The beef live**r dataset** contained two types of spectra of liver samples from healthy animals, one acquired in positive ion mode and the other in negative ion mode, both in sensitivity mode using a Synapt G2-S Q-TOF instrument (Waters, Wilmslow, United Kingdom). These large and medium datasets are used to investigate the transfer learning and representation learning approaches. SpiderMass is based on WALDI process which corresponds to MALDI with water as matrix [42]. We trained the CNNs with the MALDI dataset (rat brain dataset) and then perform classification (canine sarcoma and microorganism datasets) based on this training.
5. The human ovary dataset 1 represented two classes of serum, healthy and cancerous. High-resolution spectra are acquired via ProteinChip weak-cation-exchange interaction chips (WCX2, Ciphergen Biosystems, Inc., Fremont, CA, USA) and surface-enhanced laser-desorption/ionization (SELDI) TOF technology (QSTAR Pulsar I, Applied Biosystems, Inc., Framingham, MA, USA). This dataset has been used previously [43].
6. The human **ovary dataset 2** (as above) contained spectra acquired in low-resolution using a WCX2 protein chip via a Protein Biological System II (PBSII) SELDI-TOF instrument. This dataset has been used previously [44].

We validated our transfer and representation learning approach using these two small well-controlled, independent, and publicly available clinical MS datasets. These two datasets are based on SELDI process, we trained the CNNs with the MALDI dataset (rat brain dataset) and then perform ovarian classification based on this training. SELDI is an old ionization method which has small utility nowadays since it yields only a subset of the most abundant peptides and protein fragments. Some SELDI platforms may not be suitable for routine clinical diagnosis and struggle to prove their worth as reliable tools [45]. Low-resolution can make close species in m/z difficult to distinguish and give rise to coalesced features. Nevertheless, due to its easy-to-use quantitative screening procedures, SELDI low-resolution still can be used for general description of proteins [46]. Rather than assessing the utility of these technologies or instruments, our goal in this paper is to see the problem from the user standpoint. We illustrated the strength of our methodology as a solution to multiple real-life constraints such as the fact that the user is confronted with several types of data generated by different devices, and often in a limited size.

All classes considered in this study are non-overlapping. Datasets availability and complete descriptions are provided in the supplementary section. Table 1 lists the sample material classes used in this study.

**Table 1:**
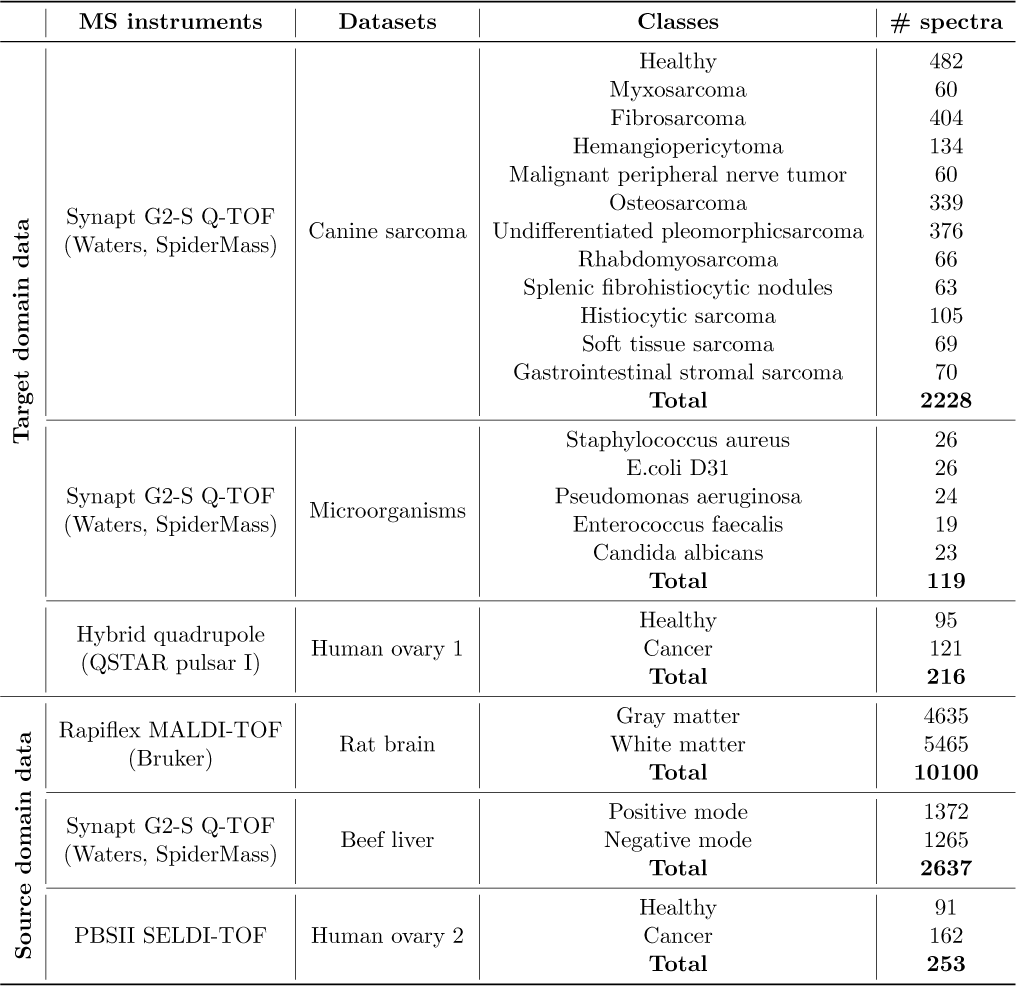
Description of datasets.

### 2.2 Hyper-parameter search

We evaluate the effects of hyper-parameter value alterations on the classification accuracy of the clinical datasets by CNNs. Contrary to some 1D data where CNN architecture and fine-tuning played a marginal role in classification performance [47]. MS data hyper-parameters selection have a huge impact on the performance, as strong as images. We begin with an investigation of the optimal convolutional filter size for the extraction of spectral features, followed by a search of various learning rate, including 0.1, 0.01, and 0.001. We reduce the learning rate when the validation set accuracy stopped improving during 10 epochs. We also investigate the use of two optimizer algorithms, including Adam and Stochastic gradient descent (SGD). We search the use of various batch sizes, including 64, 128, and 256. This evaluation is also done in terms of regularizer technique by adding either batch normalization, dropout of 0.5 or L1/L2 regularization after each convolutional layer.

### 2.3 Evaluation protocol

We import all MS datasets without undergoing any preprocessing step. We binne each dataset (see matrix construction section in the supplementary material) and scale it linearly between 0 and 1. Datasets are divided randomly into three subsets, one for training, one for validation, and one for testing with ratios of 60%, 20%, and 20%, respectively. Performance of classifiers is measured by four metrics: global accuracy (over all classes), sensitivity, specificity, and confusion matrix as an indicator on how is simple or hard for the classifier to distinguish between different classes. CNNs weights are initialized with He normal distribution since Relu/Leaky Relu is used as the activation function [48], except for the output layer, where a sigmoid function is used for binary classification or a softmax function for multi-class classification. Classification accuracy is averaged over 10 independent iterations. These subsets are computed for each iteration using a stratified sampling to maintain the original proportion of minority classes. A loss function is weighted during the training process for samples from under-represented classes in the datasets. Only the best hyper-parameters are used for the evaluation process. We use the *early stopping* to evaluate the model periodically. The model is saved only if there is an accuracy improvement of the validation set over 10 epochs and thereby use those weights for testing. The training process ends usually after about 50 epochs.

#### Protocol for evaluating prominent 2D-CNN architectures adapted to 1D input

The aim of the first experiment is to evaluate and compare the application of three prominent CNN architectures for classifying spectra in clinical datasets. The first of these is variant Lecun adapted from [49], the second is variant LeNet [26], and the third is variant VGG9 adapted from [31]. Variant Lecun (model 1) contains two convolutional layers and two fully connected layers. Variant LeNet (model 2) includes three convolutional layers and two fully connected layers. Variant VGG9 (model 3) is the deepest, with six convolutional layers and three fully connected layers. CNN architectures share the same characteristics and follow the same principles whether they are 1D or 2D. The basic difference is the dimension of the input signal and consequently how filters slide across the data. Models 1 and 3 were described in the literature and are modified slightly to fit our 1D data classification problem. Convolutional modules and pooling size are adapted to 1D input. The same number of filters is used but they are expanded to account for spectral features larger than those extracted from images (kernel size adjusted to 3×3). No zero padding is needed because all of the spectra start and end with a zero value and have the same length through binning. For model 1, two fully connected layers out of three from the original LeNet architecture are kept. The adaptation of 2D-CNN architecture to 1D input was described previously, for example in Inception modules [37] according to data specificities. Using this approach, we expect to determine what model depth and hyper-parameters are optimal for MS spectra classification. This evaluation allow assessment of layers number required for spectral feature extraction, especially in the case of highly heterogeneous biological classes such as canine sarcoma types. That is where CNNs robustness or invariance to spatial transformation proprieties handle the inter-class variability. This variability results mainly in pics translation (shift) and intensities variability from one spectra to another. Figure 1 illustrates the typical within-class variance of three spectra in the myxosarcoma tissue.

**Figure 1:**
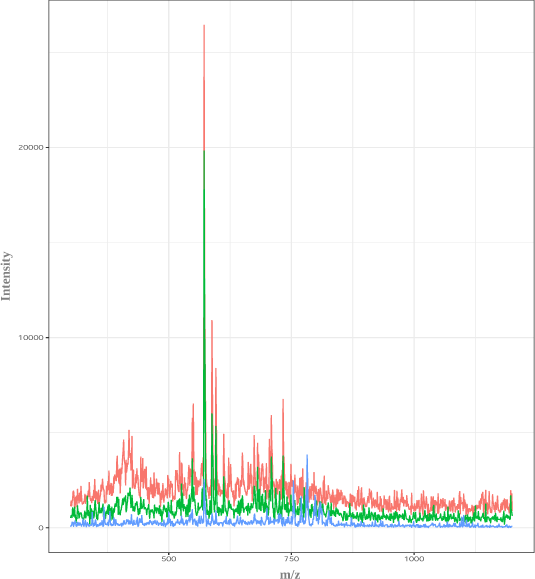
An example of three spectra from canine myxosarcoma type indicating the within-class variation

#### Protocol for evaluating transfer learning

The aim of the second experiment is to evaluate model improvement by CNNs spectral transfer learning. The three CNN architectures are trained on the large MALDI-MSI rat brain dataset with all weights initialized according to He normal distribution. Rat brain dataset is chosen as the source domain as it is the largest dataset in our study. The decision layers (fully connected layers and sigmoid layer) of the CNN networks are not useful, since the MALDI-MSI and datasets are from different domains. The representation model weights (i.e. the convolutional portion) are then frozen so that they would not be updated during back-propagation, the decision layers are removed, and the new specific decision layers dedicated to smaller clinical datasets are trained. The evaluation of transfer learning using the canine sarcoma and microorganism datasets are illustrated in the black portion of Figure 2.

**Figure 2:**
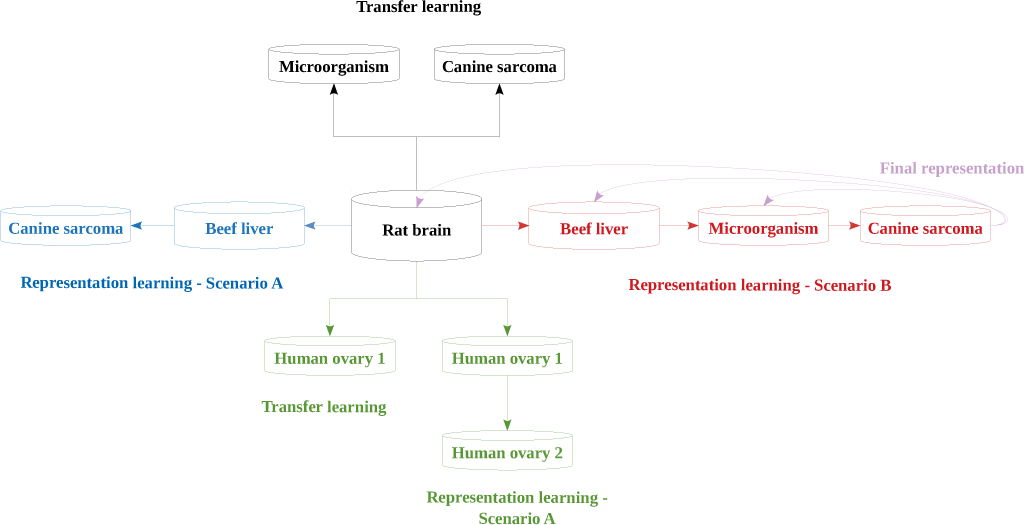
Workflow diagram of 1D-CNN classification by transfer learning and cumulative learning on clinical datasets

#### Protocol for evaluating representation learning

Transfer learning in some cases may not be enough as an aid in classifying biologically similar materials using CNN models. This proximity is reflected in a high degree of confusion between classes. This is typically the case when the biggest dataset which is supposed to be used to learn the pivotal data representation is not big enough. In addition, low-resolution or data heterogeneity can complicate more the classification task. We therefore propose two approaches to developing 1D-CNN representation (cumulative) learning:

#### Scenario A

The first step is to train CNN architectures on the MALDI-MSI rat brain dataset as described before for transfer learning. The representation model weights are then fine-tuned, the decision layers (i.e. fully connected and last sigmoid) are removed, and new decision layers are trained with the beef liver dataset. Beef liver CNN weights (i.e. data representation) are thus initialized from the rat-brain-trained CNN representations. Finally, the beef liver CNN representation weights are frozen and new specific decision layers (fully connected and softmax) are added and trained using the canine sarcoma dataset, as illustrated in the blue portion of Figure 2.

#### Scenario B

CNN architectures are trained on MALDI-MSI rat brain and fine-tuned with the beef liver dataset as described in Scenario A, but instead of testing this representation on the canine sarcoma dataset, an additional representation learning is added. Beef liver CNN representation weights are fine-tuned, decision layers (fully connected and sigmoid) are removed and new specific decision layers are added and trained using the microorganism dataset, before freezing convolutional layer weighting barring the last one and training new specific last convolutional and decision layers on the canine sarcoma dataset, as illustrated in the red portion of Figure 2.

The green portion of Figure 2 illustrates the application of transfer and cumulative learning (Scenario A) approaches to the two public ovarian datasets following the same strategy as described before.

The final representation (pink links on Figure 2) obtained from Scenario B is tested, after adapting the output space dimensionality (number of classes) and the activation function of the last fully connected layer, on rat brain, on beef liver and on microorganism spectra separately. The objective is to assess how much learning skill the final CNN representation gained or lost of MS knowledge through successive training on MS datasets.

Such an approach differs from standard transfer learning by different aspects:

- the number of output classes: because of its abundant categories and large number of images, ImageNet is used widely as the source dataset in transfer learning cases. The typical transfer learning operation consists of using a pre-trained model, for instance on 1,000 different ImageNet dataset classes and applying it to a new classification problem (possibly after fine-tuning to adapt to the new problem), which usually involves a much smaller number of classes to be predicted. In this study, the transfer learning approach comprised training a dataset with only two output categories (rat brain gray and white matter) and efficient transfer of the model to classification problems with 2, 3, 5 and even 12 output categories.
- the diversity of the target tasks: in standard transfer learning, the target tasks are similar and thus rely on similar input data features (i.e. image classification task). In this study, transfer learning and cumulative representation learning are applied to different biological contexts (i.e. diseases) that are unlikely to share features. Our results show that CNNs are powerful tools for learning generic and “potentially general” representations from spectra having no intuitive relationship to the medical target (from MALDI-MSI to intensity data) or sensitivity to acquisition instrument diversity.
- The accumulation of learning representations through several phases of representation model training up to the final decision level (fully connected layers and softmax/sigmoid layer): standard transfer approaches to learning generic representations require that the initial model be trained with as much data as possible to integrate all input data essential features. This leads to poor results with small datasets. In this study, instead of considering only transfer learning (which would be a one-shot representation model learning), the same representation model is trained for several tasks in sequence to converge on an optimal model. It thus learns cross-classification tasks and cross-instruments representation and thereby becomes capable of smoothing fluctuations in MS instruments performance, which leads to significantly improved classification accuracy.

#### Protocol for comparing our approach with other ML approaches

Some 1D-CNNs have been found superior to conventional and popular algorithms for classifying raw data [26, 36, 37]. The aim of our third experiment is to compare our 1D-CNNs to conventional ML algorithms, namely SVM, RF, and LDA. To make such a comparison valid, all spectra are binned similarly, and the same ratio of training, validation and test subsets is conserved. These conventional algorithms are not designed to classify MS spectra that have not been preprocessed. In order to compare their performance to that of CNNs on raw data, the spectra are corrected using sequential preprocessing means provided in MALDIquant package (version 1.19.3) [50]. The preprocessing comprises five steps, each of these feasible using any of several methods. For the present purpose, the most standard methods are chosen: (1) Log-intensity transformation. (2) Baseline subtraction using the statistics-sensitive non-linear iterative peak-clipping or SNIP algorithm [51]. (3) Normalization with the total ion count (TIC). (4) Alignment using a cubic warping function as described previously in [17]. We align spectra by class, namely spectra of each class are aligned separately. A non-linear cubic warping function is computed for each spectrum by fitting a local regression to the matched reference peaks. (5) Peaks are detected using the median absolute deviation. Spectra are aligned prior to peak detection in order to preserve all peak information (height, width, and spatial distribution) and thereby ensure the best alignment. For SpiderMass, the irradiation time is set at 10 sec at 10 Hz, giving an average of 10 individual spectra (1 per laser pulse) over the 10 sec period. Each microbial class was based on spectra obtained from a single acquisition. This was not the case for canine sarcoma where we merge spectra from different biopsies. Thus, the canine sarcoma and microorganism datasets have different artifacts and therefore required different preprocessing. Ovarian datasets are preprocessed following the same preprocessing strategy. The hyper-parameters for each algorithm are tuned with a grid search and are described in the supplementary section. Only the optimal hyper-parameters are used for the evaluation. Chi-square (*χ*^2^) statistic is used to reduce data dimensionality before feeding to the classification algorithms. Only features exceeding a threshold of 0.1 are selected. In addition, colinear variables are removed for LDA classification to allow the matrix inversion.

## 3 Results

### 3.1 Hyper-parameter search

The regularizer technique, the optimizer algorithm and the learning rate reveal significant effects on classification accuracy. Batch normalization, used after each convolutional layer to avoid over-fitting, is found superior to the dropout technique and L1/L2 regularization. The Adam optimizer with default hyper-parameters *β*_1_ = 0.9, *β*_2_ = 0.999 and a constant learning rate of *η* = 0.001 is found superior to the SGD algorithm. Adam is carried out using a cross-entropy loss function. We also find that Max-Pooling is very important in order to account for peak shift invariance along the m/z dimension. Since batch size do not affect the results, it is set at 256. ReLu (models 1 and 3) and Leaky Relu (model 2) are chosen as the activation function for each convolutional layer. No big discrepancies are noticed in the learning curves except for canine binary classification task because of the large class imbalance. We therefore reduce the patience epochs number from 10 to 5 for this task to avoid over-fitting. We notice that large filter size is more effective than image-optimized filtering (pixel features). This indicates that features extracted from spectral data differ from those seen in images. Kernel sizes are large enough to cover the largest peaks in the samples so that the model do not need many layers to avoid over-fitting, but not so large that detailed information is lost due to smoothing effects. CNNs architectures and their best hyper-parameters are shown in Figure 3.

**Figure 3:**
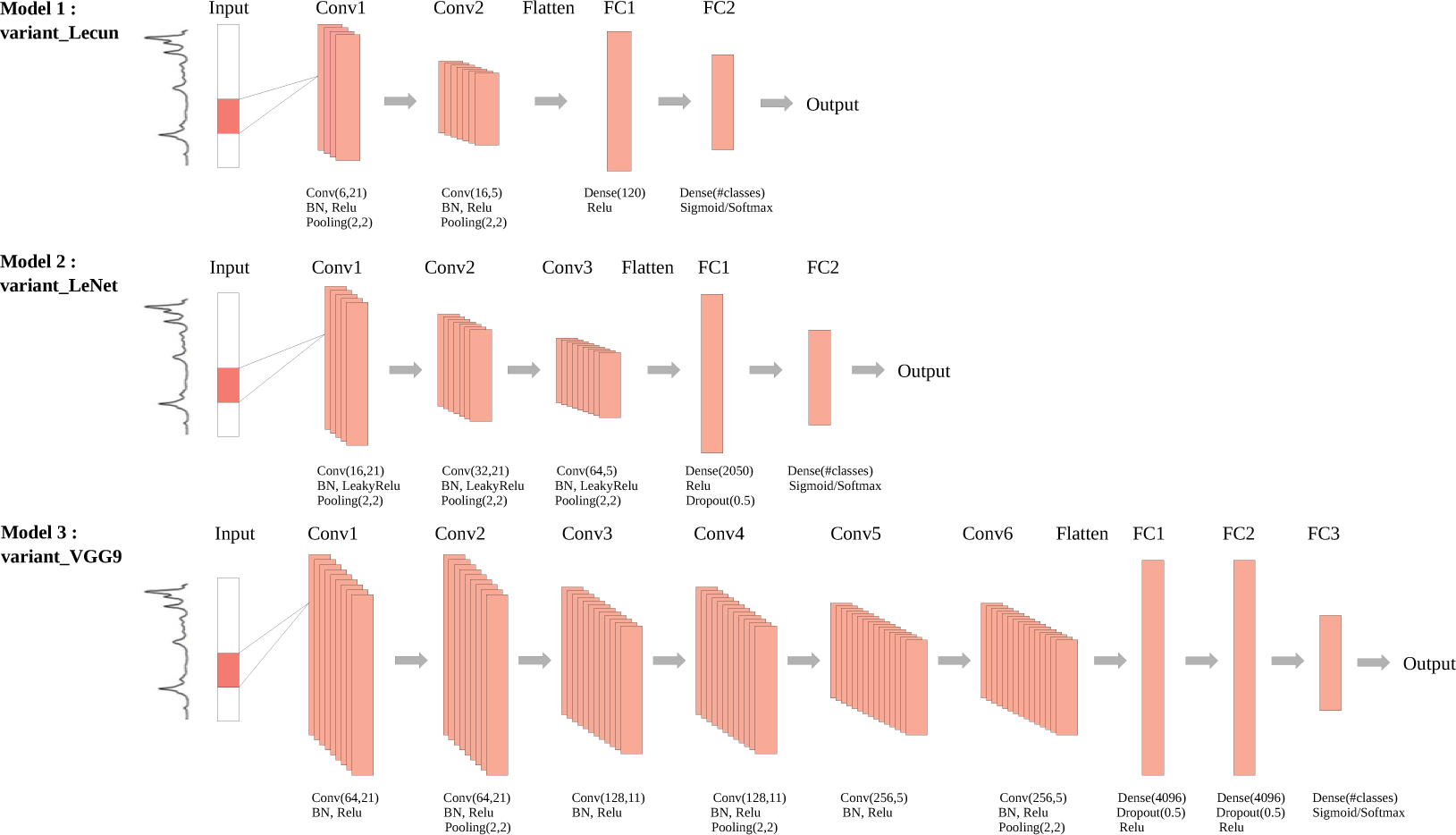
Architectures of the three CNN models. Convolutional layers are labeled as Conv, flatten layer as Flatten, and fully connected layers as FC

### 3.2 Comparison of the classification performance of three CNN architectures

The CNN architectures are compared using the cancer and microorganism datasets. In order to evaluate the effect of varying the number of CNN layers on classification performance, CNNs containing four (variant Lecun, model 1), five (variant LeNet, model 2) or nine layers (variant VGG9, model 3) are evaluated and compared. The statistical significance in classification accuracy between the first and the second best result is computed with a t-test over 10 independent iterations (p.values *<* 0.001). For the microorganism dataset, two multi-class classifications based on standard classification for clinical purposes are considered. The first is a 3-class model intended to identify the sample as yeast (*c. albicans*), Gram-positive bacteria or Gram-negative bacteria. The second model is intended to allow identification of each of the five microorganisms. For canine sarcoma classification, binary (2 classes) classification of tissues as healthy or cancerous is sought first, followed by differentiation of sarcoma type (12 classes). Table 2 lists the classification accuracy for each dataset. All sensitivity, specificity and confusion matrix metrics associated with each dataset are described in the supplementary section (Tables 1 to 8).

**Table 2:**
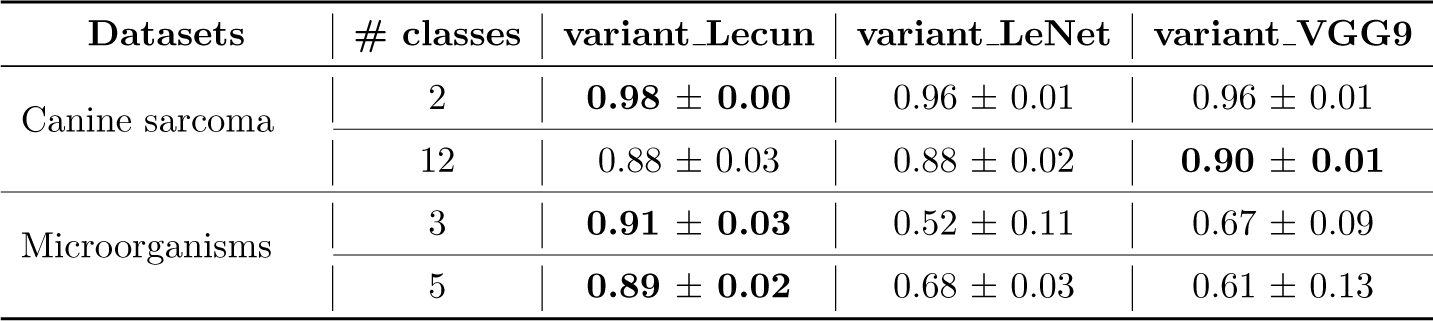
Overall accuracy of SpiderMass spectra classification using three CNN architectures. The best result for each task over 10 independent iterations is indicated in boldface

All three CNNs architectures perform poorly on 3 of the 4 tasks, which is not surprising because of the low number of spectra used for the training. Variant Lecun is the best at binary classification of canine sarcoma, but when the number of classes is expanded to 12, variant VGG9 is slightly better. This suggests that deep CNNs might be better at sorting out heterogeneous samples. Errors in the confusion matrix are distributed uniformly across classes (supplementary section Table 4). Deeper 1D-CNN architectures (16-layer and 19-layer versions of VGG) are also tested on the canine sarcoma dataset, but do not produce a better classification result (data not shown). Variant Lecun is the best at classifying microorganisms, using the 3-class or the 5-class model. Accuracy suffers quickly from over-fitting when a deep architecture such as variant LeNet and variant VGG9 are used on data of this size. The only classification that could be described as accurate is for canine sarcoma versus healthy tissue (binary classification) by variant Lecun with an average accuracy of 0.98. Based on this result, we focus our subsequent efforts on the canine sarcoma and microorganism multi-class classifications.

**Table 3:**
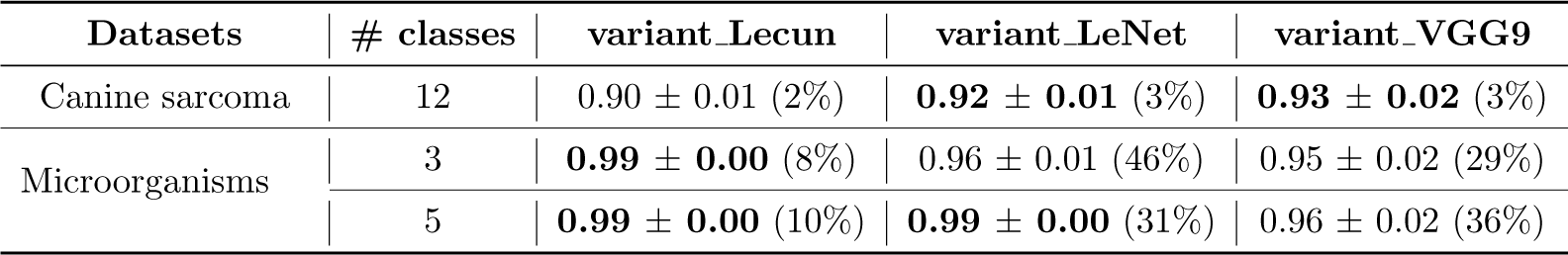
Overall accuracy of SpiderMass spectra classification using three CNN architectures after transfer learning. The improvement in performance from scratch is expressed as a percentage

**Table 4:** Overall accuracy of canine sarcoma classification by the three CNN architectures. The improvement in performance is expressed as a percentage relative to learning from scratch^∗^, to transfer learning^∗∗^, and to Scenario A^∗∗∗^

| Protocol | variant_Lecun | variant_LeNet | variant_VGG9 |
| --- | --- | --- | --- |
| Scenario A | $0.92 \pm 0.01$ (4%* 2%**) | **$0.95 \pm 0.01$** (7%* 4%**) | $0.94 \pm 0.01$ (4%* 1%**) |
| Scenario B | $0.95 \pm 0.02$ (7%* 5%** 3%***) | **$0.99 \pm 0.00$** (10%* 7%** 3%***) | $0.96 \pm 0.00$ (6%* 3%** 2%***) |

### 3.3 Transfer Learning

In order to improve the classification performance, we use CNN architectures trained on the large MALDI-MSI rat brain dataset and test them on small clinical datasets. We obtain nearly (0.99 ± 0.00) for MALDI-MSI dataset binary classification with the three CNN architectures. Transfer learning allow the model to learn and detect generic representations of MS peaks. By freezing the lower CNN levels, we assume that the model extracts the right patterns from the MALDI-MSI spectra, and that only the high level is needed to take into account specific SpiderMass peak features. As shown in Table 3, transfer learning clearly improve the accuracy of classification of both small SpiderMass datasets compared to the models trained from scratch (without transfer learning).

Gains in the accuracy of canine sarcoma differentiation are obtained for all three architectures, although improvements are still needed. Variant LeNet and variant VGG9 predict the correct classes with almost equal success, but both fail to separate some classes, as shown in the confusion matrix in supplementary Table 10. The 3-class microorganisms classification is improved somewhat for all three architectures. Improvements is considerable also for the 5-class task, and huge in the case of variant VGG9. Transfer learning by variant Lecun lead to the best performances in the experiment. These results suggest that training a CNN model with extracted spectral features transferred even from an unrelated field is better than training it with spectral features learned from scratch with a small dataset. The aim of the following experiments is to improve the canine sarcoma multi-class classification performance.

### 3.4 Cumulative Representation Learning

To improve further the accuracy of the canine sarcoma multi-classification, CNNs are trained using the large MALDI-MSI rat brain dataset and then fine-tuned using the SpiderMass datasets. Two scenarios are tested: (A) training on intermediate beef liver and then canine sarcoma dataset. We obtain nearly (0.99 ± 0.00) for rat brain and beef datasets binary classification with the three CNN architectures. Although not biologically related to the sarcoma context, beef liver recognition allows the model to appropriate the clinical data and their specific characteristics to improve its generalization capability in the second step; (B) training on beef liver, then on microorganisms and lastly on canine sarcoma dataset.

As shown in Table 4, Scenario A improves the classification accuracy considerably relative to learning from scratch and slightly relative to transfer learning, the best improvements is obtained for variant LeNet. Scenario B provides a slight additional improvement over Scenario A, and the greatest accuracy is achieved also with variant LeNet architecture. The effectiveness of the cumulative knowledge method is thus apparent, enabling the CNNs to distinguish not only cancerous versus healthy tissues (binary classification), but also the different cancer types (see confusion matrices in Table 16 for Scenario A and in Table 18 for Scenario B, supplementary section) despite the large number of classes, the small size and the heterogeneity of the dataset. We test CNNs configured with different numbers of frozen layers using transfer and cumulative representation learning in order to evaluate the trade-off between freezing and fine-tuning. Freezing all convolutional layers (i.e. the representation portion) and re-training all fully connected layers (i.e. the decisional portion) gives a configuration that outperform the others. Except for Scenario B where the best architecture is obtained by freezing all convolutional layers barring the last one. We test the same protocols on datasets with a smallest bin (binned at 1 instead of 0.1), similar improvements in accuracy are observed, except that the variant VGG9 architecture outperforms other networks. This may suggest that an architecture with 3 convolutional network may be sufficient with data binned at 0.1, while a deep architecture such as variant VGG9 may be needed in case of too compressed data.

Cumulative learning strategy brings new questions: how generalizable is the final representation after several steps of cumulative learning? Is the final representation more specifically adapted to the last dataset used to accumulate MS knowledge? Let us first remind that the classification accuracy obtained by CNNs from scratch on data used for training (rat brain and beef liver) and after transfer learning for microorganisms (Table 3) is equal to 0.99. Testing the final cumulative representation of variant LeNet (pink links from Scenario B) on rat brain, beef liver and microorganism datasets separately preserved a classification accuracy of 0.99. This indicates that the CNN model accumulates MS knowledge through the successive training phases without any loss of generalization. It suggests that a “generic” representation of MS data for classification tasks might exist and that the resulting cumulative representation is robust to the organism, to the tissue phenotype, and to the instrument variability.

#### Public MS datasets

The classification accuracy on the two ovarian datasets were compared previously to explain how the choice of the MS instrument, its resolution or preprocessing steps become an obstacle to the reproducibility and reliability of patterns. The aim of this experiment is not to compare our results to the reported results in the original paper analyzing these datasets, first because the commercial Proteome Quest software is used for classification, and then the preprocessing strategy is different in the original paper (refer to [52] for more details). Our purpose is to demonstrate the efficiency of our learning methodology capable of handling multiple MS features : ionization sources (from WALDI to SELDI), resolutions (from high to low), and mass ranges (from lipids to proteins). We assess CNNs performance using the same training and evaluation approach. Only with variant LeNet architecture, because of its superior performance with SpiderMass datasets and its low computational resources needed. Variant LeNet is thus trained on the rat brain dataset as the source domain, followed by the transfer learning protocol using the high-resolution dataset and representation learning (Scenario A) using the low-resolution dataset.

Transfer learning improve classification accuracy from 0.78 for training from scratch to 0.98 for the high-resolution dataset (Table 5). With the low-resolution dataset, accuracy is improved from 0.80 to 0.83 by transfer learning and up to 0.99 by cumulative learning. These results show that in contrast with the previously reported lack of sensitivity and specificity of low-resolution MS datasets for diagnosis and the fact that the identified fragments usually belong to nonspecific proteins unlikely to be directly associated with the disease [45]. Our CNN representation model allow very high classification accuracy, without the need for spectral preprocessing steps, and while the model is trained on a dataset with different MS features.

**Table 5:** Overall accuracy of a variant LeNet architecture at classifying ovarian cancer serums; percent improvement relative to learning from scratch^∗^ and to transfer learning^∗∗^

| Dataset | # classes | variant_LeNet | Transfer learning |
| --- | --- | --- | --- |
| Human ovary 1 | 2 | 0.78 ± 0.02 | **0.98 ± 0.00** (24%*) |

| Dataset | # classes | variant_LeNet | Transfer learning | Cumulative representation learning |
| --- | --- | --- | --- | --- |
| Human ovary 2 | 2 | 0.80 ± 0.00 | 0.83 ± 0.02 (3%*) | **0.99 ± 0.00** (24%* 19%**) |

### 3.5 Comparison of our 1D-CNN against other ML approaches

Spectra are corrected using sequential preprocessing of five steps: Log-intensity transformation, baseline subtraction using SNIP algorithm, TIC normalization, alignment using a cubic warping function, and peaks detection using the median absolute deviation. Alignment step consists in identifying landmark peaks also called reference or anchor peaks. Reference peaks are found in each of the microbial classes. However, some canine sarcoma classes are obtained from different biopsies (dogs of different breed and age) with signal acquisition spread over days. These resulted in a high spectral variability, which made reference peak identification difficult. The ideal alignment would be based on reference peaks occurring in all spectra of each class. When common peaks are absent or fewer than our threshold of four peaks (e.g. for some canine sarcoma classes in binary and multi-class classifications), peaks occurring in half of the spectra are used. These might not be biologically relevant and might had a negative impact on classification performance. Such choices are constraining and depend on the specialist’s expertise, as mentioned in the introduction section. The performance of the best 1D-CNN, that is, of all models and approaches combined, from scratch for binary canine sarcoma classification, after transfer learning for microorganisms and human ovary 1, after cumulative learning (Scenario A) for human ovary 2 classification, and Scenario B for multi-class canine sarcoma are compared to SVM, RF, and LDA (Table 6).

**Table 6:** Overall accuracies of clinical spectra classifications by SVM, RF, and LDA; percent difference to 1D-CNN trained from scratch^∗^, after transfer learning^∗∗^, and after representation learning^∗∗∗^

| Datasets | # classes | SVM | RF | LDA |
| --- | --- | --- | --- | --- |
| Canine sarcoma | 2 | $0.76 \pm 0.16$ (22%*) | **$0.96 \pm 0.01$** (2%*) | $0.88 \pm 0.17$ (10%*) |
| | 12 | $0.52 \pm 0.19$ (47%***) | **$0.65 \pm 0.01$** (34%***) | $0.61 \pm 0.02$ (38%***) |
| Microorganisms | 3 | $0.87 \pm 0.02$ (12%**) | **$0.95 \pm 0.02$** (9%**) | $0.87 \pm 0.01$ (12%**) |
| | 5 | $0.54 \pm 0.35$ (45%**) | **$0.86 \pm 0.01$** (13%**) | $0.51 \pm 0.26$ (48%**) |
| Human ovary 1 | 2 | $0.66 \pm 0.24$ (49%***) | **$0.91 \pm 0.02$** (8%***) | $0.85 \pm 0.06$ (15%***) |
| Human ovary 2 | 2 | $0.60 \pm 0.05$ (65%**) | $0.88 \pm 0.03$ (12%**) | **$0.97 \pm 0.00$** (2%**) |

RF gives the best result and outperforms the two other methods in SpiderMass and human ovary 1 datasets classification, especially in canine binary task, where the classification is the more accurate. LDA outperforms the two other methods for the human ovary 2 dataset classification. The computing time of conventional algorithm is much higher than CNNs, first because the Scikit-learn library does not support GPU-based computing, and also because a few more minutes are needed to carry out the necessary preprocessing steps. In addition, some inter-spectra steps such as normalization and alignment are applied only when the acquisition process is completed. While our pre-trained CNN models can be applied even during spectra acquisition since each test spectra could be analyzed separately.

## 4 Discussion

CNNs have become common tools in several research areas. They are designed to extract spatial features from input signals with different levels of abstraction. Many challenges remain fully exploiting CNNs on biomedical data, owing to data high-dimensionality, heterogeneity, and irregularity. Following their success in computer vision, the first results of deep learning methods to clinical data are obtained on clinical imaging (e.g. classification, segmentation, etc.). Medical images are different from ImageNet object scenes, persons, and plants, among others. Nevertheless [53–56] demonstrated that we could classify and predict outcomes from medical images using a CNN model trained on ImageNet. Authors show that the features extracted from the ImageNet database are generalizable and can be applied to alternative tasks and datasets. Our paper was inspired by these efforts on transfer learning, transferring the representation learnt on one dataset to another that intuitively do not seem to share common features, but goes beyond by accumulating the knowledge of MS data space through learning a representation on several datasets, more than two and more diverse too. We have investigated here the performance of CNNs in the classification of 1D mass spectra generated for a variety of classification purposes. This study shows for the first time the use of representation learning for spectrum classification using MALDI-MSI and other types of datasets generated in vastly different biological contexts, on different organisms, acquired by a variety of instruments, with a variety of MS ionization technologies, in different mass ranges at different resolutions, and with or without chromatographic phase. Our extracted MS representation was designed by accumulating mass spectral knowledge through multiple training steps on small datasets. It was effective even on low-resolution data despite its limited potential in precision proteomics. One would believe that the effectiveness of transfer learning depends on the relationship between source and target domains. However, our results showed the opposite even with poorly related domains, the performance of the final model increased. In addition, our strategy provided a viable alternative when transfer learning was inadequate, as was the case for low-resolution, heterogeneous data, or when the source domain dataset was not large enough. The particularly interesting novelty here was that the model can be pre-trained on a dataset containing only two output categories and yet predict 2, 3, 5 and even 12 outputs, unlike what has been described in the literature. The choice of our learning order in this paper was based first on the data size, then on the classification accuracy, and finally on the data resolution. Researchers can subsequently adapt the learning order according to the specificity and availability of their data.

It is well known that the success of CNNs is strongly dependent on the amount of data available for its training. To overcome the limitations inherent in small numbers of training samples, we tested dataset augmentation. To the best of our knowledge, biological 1D data augmentation has not been described elsewhere, and this may be because it is not sufficient to reproduce technical variability by adding noise plus baseline and peak misalignment. Biological variability (difference between individuals) must also be introduced, in the form of biologically relevant peak presence/absence and intensity changes. All classification accuracies on augmented SpiderMass data were below 0.60 (results not shown). This indicates that better understanding of biological variability is still required in order to deal with data augmentation and increase the number of samples without compromising biological information.

The CNN model was able to classify raw MS data without preprocessing steps, not even removal of peaks corresponding to molecule isotopes in SpiderMass datasets, thus bypassing the expert parameter setting step. This performance capability was due to convolutional filters that allow CNN architecture to learn peak patterns rather than only considering each m/z intensity value separately as do conventional ML algorithms. More importantly, significant variations of the overall signal intensity due to biological heterogeneity (not all peaks showing up in each sample) and non-reproducible technical factors (noise and baseline distortion) can be filtered by CNNs to increase the robustness of molecular pattern recognition. Combined Max-Pooling and convolution filters allowed the model to ignore peak misalignment, thus reducing the need for data preprocessing before classification. Such end-to-end trainable systems that work with raw data offer a superior alternative to pipelines in which each step is trained independently or handcrafted to find the best combination of parameters. Interrogation of pre-built models will be fast and can be implemented in the SpiderMass system to provide real-time feedback to the end user.

## 5 Conclusions and Future Works

In this work, we have investigated the potential of using a pre-trained 1D-CNN for analyzing clinical datasets drawn from vastly different clinical situations: cancers diagnosis and microorganism identifications. Our results provide evidence that cumulative learning offers practical means of analyzing mass spectra obtained in real-world settings where the size of the dataset available for training a classification algorithm is limited. Our cumulative optimization of CNN models appeared to be better adapted than conventional ML models for mass spectra classification, even when the tasks required analysis of heterogeneous, low-resolution datasets containing several classes. In addition, cumulative CNNs appear to offer a unified solution for classification regardless of day-to-day, sample-to-sample and machine-to-machine variance. Furthermore, cumulative CNNs reduce the need to preprocess data before classification or to be concerned with noise, variability and high data dimensionality. Using raw MS data directly has the potential to contribute significantly to the development of a diagnosis workflow for rapid, efficient and reliable detection of cancers or infections. It thus opens the door to real-time decision-aid tools in clinical settings independently of the MS instrument in use. Although we have focused our study on mass spectra, we believe that the method should be applicable to other types of analyses such as Raman Spectroscopy or Nuclear Magnetic Resonance classification tasks.

In the present study, we have investigated the performance of our learning approach for MS data classification. It would be interesting to extend the investigation to analyze which data characteristics were transferred between datasets. The focus of our future research will be the interpretation of classification results in order to identify regions of interest in spectra. Successful transfer learning between lipids and proteins species suggests that CNNs have a more complex functioning than the simple identification of specific-phenotype peaks. Discriminating regions during the CNN classification may be explained by a complex spatial pattern recognition and by the ability of the model to generalize from one classification task to others even slightly related. It would be interesting to understand how such markers influence the results and to study their global dynamics of expression.

## Author contributions

K.S. has done all the experiments. F.P. and K.S. have worked on designing the studies and have written the paper. P.S. has generated SpiderMass data. N.O. provided the MALDI data. N.O. and M.W. have participated to the interpretation of the results. A.D., I.F. and M.S. have obtained funds for the project. I.F. and A.D. supervised the study. All authors have approved the final version of the manuscript.

## Acknowledgments

The operation Graham supercomputer is funded by the Canada Foundation for Innovation (CFI), the ministére de l’Économie, de la science et de l’innovation du Québec (MESI) and the Fonds de recherche du Québec - Nature et technologies (FRQ-NT). This research is supported by funding from Ministere de l’Enseignement Supérieur, de la Recherche et de l’Innovation (MESRI), Institut National de la Santé et de la Recherche Médicale (Inserm), Agence Nationale de la Recherche (ANR), SATT Nord, Institut Nationale du Cancer (INCA), and Université de Lille, with the support of “Service de coopération et d’action culturelle du Consulat général de France à Québec”

## Supplementary material

### Data availability and Tissue Preparation

1. Canine sarcoma dataset: Freshly removed tissue is snap-frozen in liquid nitrogen and stored at −80°C. Tissue samples are warmed to −20°C, sliced and then analyzed using the SpiderMass system as described previously [40].
2. Microorganism dataset: Gram negative (*E. coli D31, Pseudomonas aeruginosa*); Gram positive (*Staphylococcus aureus, Enterococcus faecalis*) and Yeast (*Candida Albicans*) are cultivated in poor broth medium as described previously [41].
3. Rat brain dataset: The Wistar rat (Ratus norvegicus) are sacrificed after behavioral practical in the University of Lille, their brains are collected and snap frozen in liquid nitrogen and stored at −80°C. There are then sectioned at 12µm using a cryostat (Leica microsystems) and thaw-mounted on Indium Thin Oxide slides (LaserBio Labs). The 2,5-Dihydroxybenzoic acid (DHB) matrix (Sigma-Aldrich) is sublimed onto the tissue section at 150°C for 12 min using a “home-built’ sublimation device. The image is acquired at 50µm x 50µm spatial resolution in positive ion reflectron mode.
4. Beef liver dataset: Raw commercial product is sliced to suitable thickness, snap-frozen in liquid nitrogen and stored at −80°C. Tissue is warmed to RT prior to SpiderMass spectral acquisition. The dataset is generated while running a time-course reproducibility experiment in both positive and negative ionization modes.
5. Human ovarian datasets: Data sources and complete descriptions can be accessed through FDA-NCI Clinical Proteomics at https://home.ccr.cancer.gov/ncifdaproteomics/ppatterns.asp.

The datasets generated during and/or analyzed during the current study are available from the corresponding author upon request.

### Matrix construction

In this study, we focused on lipids and metabolites as the main species observed in the 100-1.600 mass/charge (m/z) range with the SpiderMass. Multiple studies have shed light on the role of lipid metabolism deregulation in cancer development [57–59]. Some tumors exhibit a lipogenic phenotype, since membrane lipids are synthesized rapidly and with high turnover in malignant cells [60, 61], suggesting the relevance of a lipid-based classification model. Recent microbial taxonomy studies have also demonstrated the possibility of biotyping pathogens using their lipid composition [62]. The classification models obtained using SpiderMass datasets are therefore based on lipid profiles. To allow fair comparison with the original paper, mass spectra acquired from public ovarian datasets (high and low-resolution) are restricted to the m/z range from 700 to 12.000 [52]. The classification models obtained using public ovarian datasets are therefore based on lipids and proteins patterns.

Raw SpiderMass spectra are converted into mzXML format using the 64-bit MSConvert tool version 3.0, part of the ProteoWizard suite [63]. Spectra with a total ion current (TIC) exceeding 1.*e*^4^ count for irradiation detection are selected using the MSnbase package (version 1.20.7, R version 3.4.4) [64]. Raw ovarian datasets are imported into a csv file format. A simple and popular method of creating an intensity matrix from multiple spectra prior to classification is spectral bucketing or binning [65]. Easy to use on MS data, binning consists of projecting spectra into “buckets” having a fixed size. All SpiderMass spectra therefore are binned to 0.1 Da for the subsequent analyses. This binning condenses canine sarcoma, microorganism, and beef liver data points respectively from approximately 119.200, 67.200, and 83.700 to 15.000 features. Rat brain MALDI spectra (m/z between 300-1.300 range) are binned to have the same dimensions as SpiderMass datasets (15.000 features) in order to allow transfer learning, which requires data of equal dimensions. To allow valid comparison with the original paper, both public ovarian datasets (high and low-resolution) are binned to m/z 7.084 as described previously [52]. Rat brain MALDI spectra are binned in this case at the same dimension (7.084) to allow transfer learning and representation learning.

### 1D-MS Data augmentation

To increase the robustness of the training and compensate for the limited number of SpiderMass spectra, the training set size is increased using the following data augmentation procedure: (1) Random noise proportional to spectrum acquisition order is added. (2) For misalignment, the shift scale is first assessed using a cubic warping function; spectra are then augmented by shifting each one along the m/z dimension using a third-order polynomial randomly between −1 and 1 Da. (3) Peak intensity values are increased by a random value ranging from 0 to 2 to produce intensity variation and peak absence/presence.

### Computing environment

The proposed 1D-CNNs are computed with Keras library ([66] version 2.2.4) on two NVIDIA P100 Pascal GPUs of 12 GB HBM2 memory (Graham supercomputer from Compute Canada at Waterloo university). Classification models with conventional ML algorithms are implemented using the Scikit-Learn library [67] (version 0.19.1) are computed on a local CPU server. All classification models are executed in a Linux environment using the Python language (Python version 3.6.8). The computational load and memory requirements for transfer and cumulative learning are low. Indeed, training time on the rat brain dataset is about 25 minutes for variant Lecun, 50 minutes for variant LeNet, and 3 hours for variant VGG. Fine-tuning on clinical datasets is completed in about 20 minute since the weighting coefficients are already determined, and all three architectures processed each test spectrum in less than one millisecond.

### Comparison of the classification performance of three CNN architectures

**Table 1:** Overall accuracy, sensitivity, and specificity values for 2-class canine sarcoma classification.

| Model 1 |  | Model 2 |  | Model 3 |  |
| --- | --- | --- | --- | --- | --- |
| Accuracy = $0.98 \pm 0.00$ | | Accuracy = $0.96 \pm 0.01$ | | Accuracy = $0.96 \pm 0.01$ | |
| Sensitivity | Specificity | Sensitivity | Specificity | Sensitivity | Specificity |
| 0.979 | 0.997 | 0.958 | 0.996 | 0.936 | 0.991 |

**Table 2:**
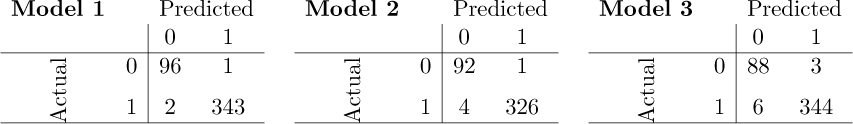
Confusion matrix for 2-class canine sarcoma classification.

**Table 3:**
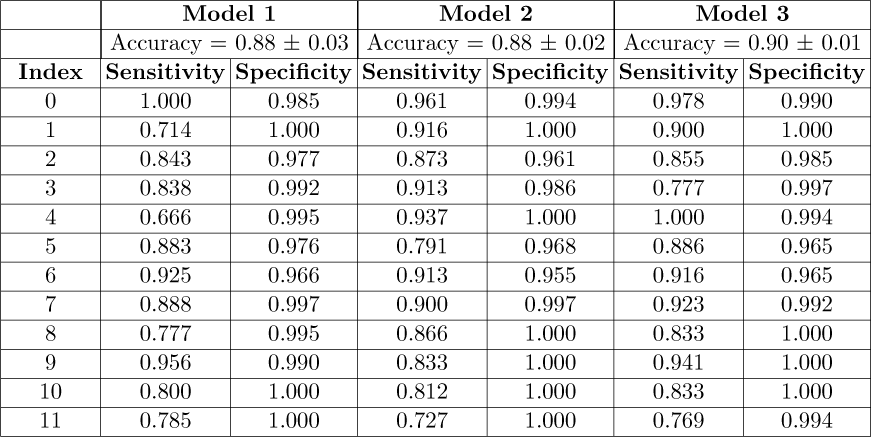
Overall accuracy, sensitivity, and specificity for 12-class canine sarcoma classification.

**Table 4:**
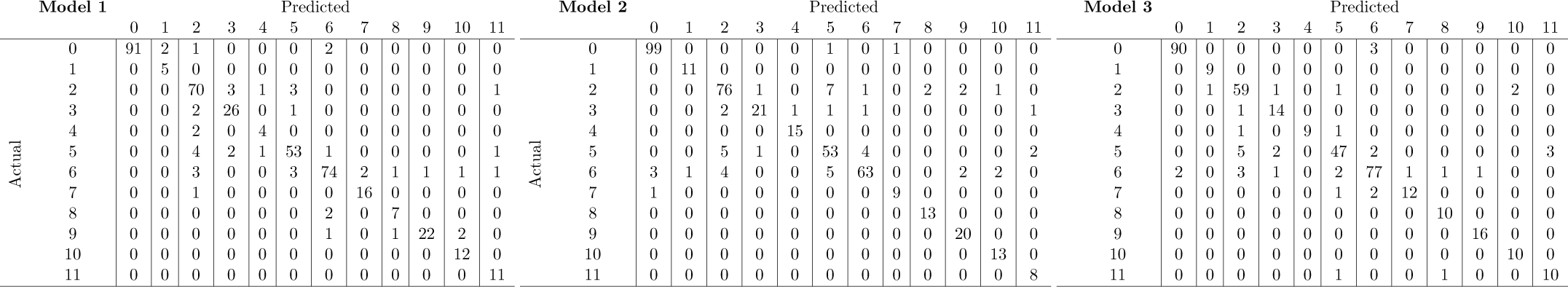
Confusion matrix for 12-class canine sarcoma classification.

**Table 5:** Overall accuracy, sensitivity, and specificity values for 3-class microorganisms classification.

|  | Model 1 |  | Model 2 |  | Model 3 |  |
| --- | --- | --- | --- | --- | --- | --- |
| | Accuracy = $0.91 \pm 0.03$ | | Accuracy = $0.52 \pm 0.11$ | | Accuracy = $0.67 \pm 0.09$ | |
| Index | Sensitivity | Specificity | Sensitivity | Specificity | Sensitivity | Specificity |
| 0 | 1.000 | 0.913 | 1.000 | 0.200 | 1.000 | 0.750 |
| 1 | 0.866 | 1.000 | 0.000 | 1.000 | 0.000 | 1.000 |
| 2 | 1.000 | 1.000 | 1.000 | 1.000 | 1.000 | 0.761 |

**Table 6:**
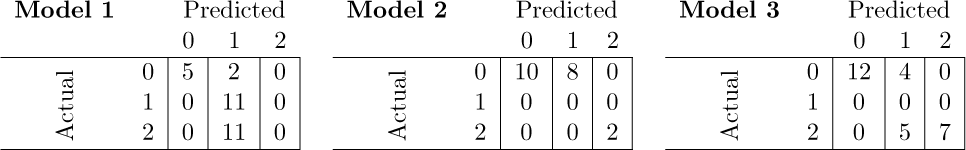
Confusion matrix for 3-class microorganisms classification.

**Table 7:** Overall accuracy, sensitivity, and specificity values for 5-class microorganisms classification.

|  | Model 1 |  | Model 2 |  | Model 3 |  |
| --- | --- | --- | --- | --- | --- | --- |
| | Accuracy = $0.89 \pm 0.02$ | | Accuracy = $0.68 \pm 0.03$ | | Accuracy = $0.61 \pm 0.13$ | |
| Index | Sensitivity | Specificity | Sensitivity | Specificity | Sensitivity | Specificity |
| 0 | 1.000 | 1.000 | 1.000 | 1.000 | 0.000 | 1.000 |
| 1 | 0.000 | 1.000 | 0.000 | 1.000 | 0.428 | 1.000 |
| 2 | 1.000 | 0.678 | 1.000 | 0.673 | 1.000 | 1.000 |
| 3 | 0.000 | 1.000 | 0.000 | 1.000 | 1.000 | 1.000 |
| 4 | 1.000 | 0.954 | 1.000 | 0.959 | 1.000 | 0.555 |

**Table 8:**
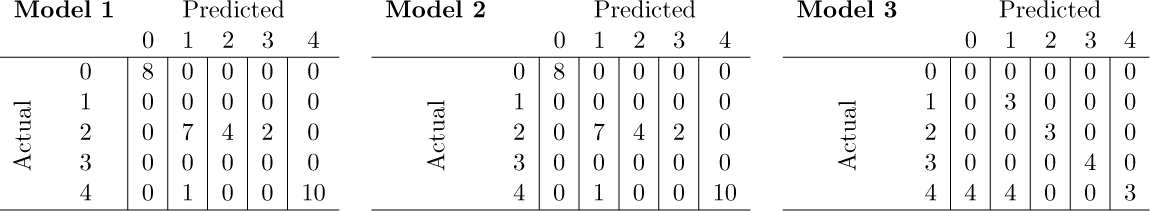
Confusion matrix for 5-class microorganisms classification.

### Transfer Learning

**Table 9:** Overall accuracy, sensitivity, and specificity for 12-class canine sarcoma classification.

|  | Model 1 |  | Model 2 |  | Model 3 |  |
| --- | --- | --- | --- | --- | --- | --- |
| | Accuracy = $0.90 \pm 0.01$ | | Accuracy = $0.92 \pm 0.01$ | | Accuracy = $0.93 \pm 0.02$ | |
| Index | Sensitivity | Specificity | Sensitivity | Specificity | Sensitivity | Specificity |
| 0 | 0.889 | 1.000 | 0.730 | 1.000 | 0.946 | 1.000 |
| 1 | 1.000 | 1.000 | 1.000 | 1.000 | 1.000 | 1.000 |
| 2 | 0.942 | 0.983 | 0.988 | 1.000 | 0.952 | 0.989 |
| 3 | 0.813 | 0.996 | 1.000 | 0.997 | 0.956 | 0.996 |
| 4 | 0.882 | 1.000 | 1.000 | 1.000 | 0.875 | 1.000 |
| 5 | 0.950 | 0.977 | 0.985 | 0.997 | 0.925 | 0.986 |
| 6 | 0.947 | 0.944 | 1.000 | 0.919 | 0.879 | 0.982 |
| 7 | 1.000 | 0.998 | 1.000 | 1.000 | 1.000 | 1.000 |
| 8 | 0.823 | 1.000 | 0.818 | 1.000 | 0.900 | 0.997 |
| 9 | 0.757 | 0.992 | 0.958 | 0.997 | 1.000 | 0.972 |
| 10 | 1.000 | 1.000 | 1.000 | 1.000 | 0.818 | 1.000 |
| 11 | 1.000 | 1.000 | 1.000 | 1.000 | 0.909 | 0.997 |

**Table 10:**
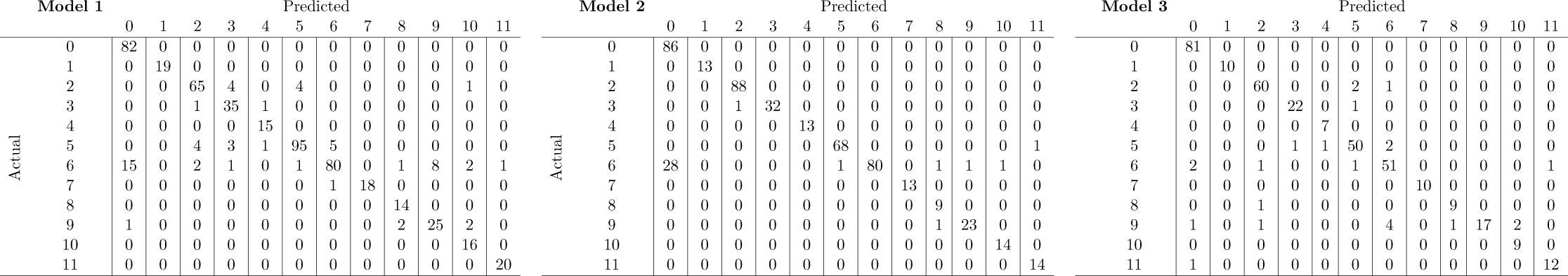
Confusion matrix for 12-class canine sarcoma classification.

**Table 11:** Overall accuracy, sensitivity, and specificity values for 3-class microorganisms classification.

|  | Model 1 |  | Model 2 |  | Model 3 |  |
| --- | --- | --- | --- | --- | --- | --- |
| | Accuracy = $0.99 \pm 0.00$ | | Accuracy = $0.96 \pm 0.01$ | | Accuracy = $0.95 \pm 0.02$ | |
| Index | Sensitivity | Specificity | Sensitivity | Specificity | Sensitivity | Specificity |
| 0 | 1.000 | 1.000 | 0.900 | 1.000 | 1.000 | 0.923 |
| 1 | 1.000 | 1.000 | 1.000 | 0.937 | 0.888 | 1.000 |
| 2 | 1.000 | 1.000 | 1.000 | 1.000 | 1.000 | 1.000 |

**Table 12:** Confusion matrix for 3-class microorganisms classification.

| Model 1 |  | Predicted |  |  | Model 2 |  | Predicted |  |  | Model 3 |  | Predicted |  |  |
| --- | --- | --- | --- | --- | --- | --- | --- | --- | --- | --- | --- | --- | --- | --- |
|  |  | 0 | 1 | 2 |  |  | 0 | 1 | 2 |  |  | 0 | 1 | 2 |
| Actual | 0 | 12 | 0 | 0 | Actual | 0 | 9 | 0 | 0 | Actual | 0 | 8 | 1 | 0 |
|  | 1 | 0 | 10 | 0 |  | 1 | 1 | 9 | 0 |  | 1 | 0 | 8 | 0 |
|  | 2 | 0 | 0 | 5 |  | 2 | 0 | 0 | 6 |  | 2 | 0 | 0 | 4 |

**Table 13:** Overall accuracy, sensitivity, and specificity for 5-class microorganisms classification.

|  | Model 1 |  | Model 2 |  | Model 3 |  |
| --- | --- | --- | --- | --- | --- | --- |
| | Accuracy = $0.99 \pm 0.00$ | | Accuracy = $0.99 \pm 0.00$ | | Accuracy = $0.96 \pm 0.02$ | |
| Index | Sensitivity | Specificity | Sensitivity | Specificity | Sensitivity | Specificity |
| 0 | 1.000 | 0.996 | 1.000 | 0.986 | 0.987 | 0.998 |
| 1 | 1.000 | 0.996 | 0.988 | 1.000 | 1.000 | 0.763 |
| 2 | 1.000 | 0.998 | 0.993 | 1.000 | 0.994 | 1.000 |
| 3 | 0.993 | 1.000 | 1.000 | 0.998 | 1.000 | 1.000 |
| 4 | 0.990 | 1.000 | 0.992 | 1.000 | 0.000 | 1.000 |

**Table 14:** Confusion matrix for 5-class microorganisms classification.

| Model 1 |  | Predicted |  |  |  |  | Model 2 |  | Predicted |  |  |  |  | Model 3 |  | Predicted |  |  |  |  |
| --- | --- | --- | --- | --- | --- | --- | --- | --- | --- | --- | --- | --- | --- | --- | --- | --- | --- | --- | --- | --- |
|  |  | 0 | 1 | 2 | 3 | 4 |  |  | 0 | 1 | 2 | 3 | 4 |  |  | 0 | 1 | 2 | 3 | 4 |
| Actual | 0 | 6 | 0 | 0 | 0 | 0 | Actual | 0 | 6 | 0 | 0 | 0 | 0 | Actual | 0 | 7 | 0 | 0 | 0 | 0 |
|  | 1 | 0 | 6 | 0 | 0 | 0 |  | 1 | 0 | 6 | 0 | 0 | 0 |  | 1 | 0 | 5 | 0 | 0 | 0 |
|  | 2 | 0 | 0 | 7 | 0 | 0 |  | 2 | 0 | 0 | 7 | 0 | 0 |  | 2 | 0 | 0 | 6 | 1 | 0 |
|  | 3 | 0 | 0 | 0 | 5 | 0 |  | 3 | 0 | 0 | 0 | 5 | 0 |  | 3 | 0 | 0 | 0 | 4 | 0 |
|  | 4 | 0 | 0 | 0 | 0 | 6 |  | 4 | 0 | 0 | 0 | 0 | 6 |  | 4 | 0 | 0 | 0 | 0 | 3 |

### Cumulative Representation Learning

#### Scenario A

**Table 15:** Overall accuracy, sensitivity, and specificity for 12-class canine sarcoma classification.

|  | Model 1 |  | Model 2 |  | Model 3 |  |
| --- | --- | --- | --- | --- | --- | --- |
| | Accuracy = $0.92 \pm 0.01$ | | Accuracy = $0.95 \pm 0.01$ | | Accuracy = $0.94 \pm 0.01$ | |
| Index | Sensitivity | Specificity | Sensitivity | Specificity | Sensitivity | Specificity |
| 0 | 0.938 | 0.938 | 0.989 | 0.996 | 0.846 | 0.994 |
| 1 | 1.000 | 1.000 | 0.916 | 1.000 | 1.000 | 1.000 |
| 2 | 0.975 | 0.978 | 0.944 | 0.991 | 1.000 | 1.000 |
| 3 | 0.964 | 1.000 | 0.916 | 0.992 | 1.000 | 0.997 |
| 4 | 0.909 | 1.000 | 1.000 | 0.997 | 1.000 | 1.000 |
| 5 | 0.968 | 0.994 | 0.968 | 0.988 | 0.972 | 1.000 |
| 6 | 0.776 | 0.989 | 0.942 | 0.985 | 0.987 | 0.939 |
| 7 | 0.833 | 1.000 | 0.916 | 1.000 | 0.933 | 1.000 |
| 8 | 1.000 | 1.000 | 0.833 | 1.000 | 0.928 | 1.000 |
| 9 | 0.958 | 0.976 | 0.952 | 0.992 | 0.739 | 0.997 |
| 10 | 0.928 | 1.000 | 0.846 | 1.000 | 0.933 | 1.000 |
| 11 | 0.857 | 1.000 | 1.000 | 1.000 | 1.000 | 1.000 |

**Table 16:**
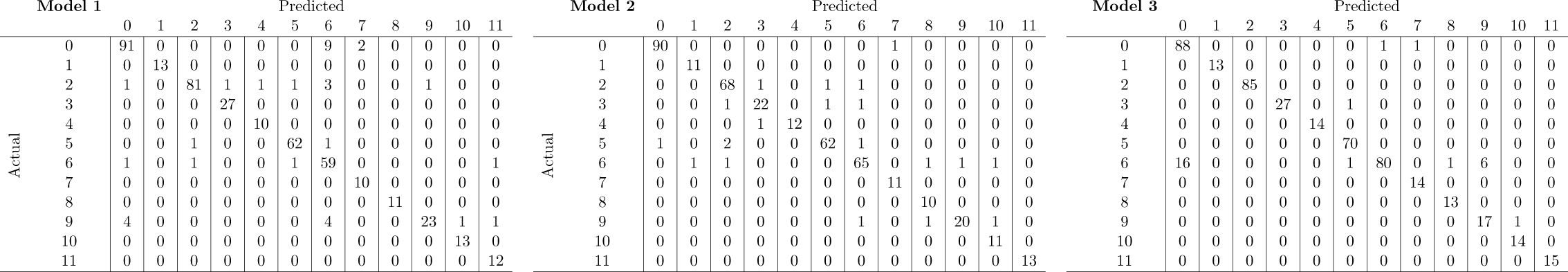
Confusion matrix for 12-class canine sarcoma classification.

#### Scenario B

**Table 17:**
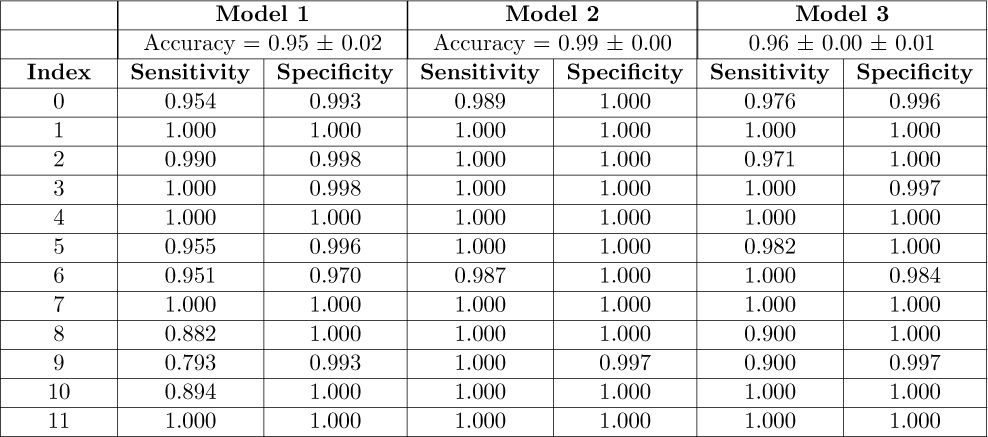
Overall accuracy, sensitivity, and specificity for 12-class canine sarcoma classification.

|  | Model 1 |  | Model 2 |  | Model 3 |  |
| --- | --- | --- | --- | --- | --- | --- |
| | Accuracy = $0.95 \pm 0.02$ | | Accuracy = $0.99 \pm 0.00$ | | Accuracy = $0.96 \pm 0.00 \pm 0.01$ | |
| Index | Sensitivity | Specificity | Sensitivity | Specificity | Sensitivity | Specificity |
| 0 | 0.954 | 0.993 | 0.989 | 1.000 | 0.976 | 0.996 |
| 1 | 1.000 | 1.000 | 1.000 | 1.000 | 1.000 | 1.000 |
| 2 | 0.990 | 0.998 | 1.000 | 1.000 | 0.971 | 1.000 |
| 3 | 1.000 | 0.998 | 1.000 | 1.000 | 1.000 | 0.997 |
| 4 | 1.000 | 1.000 | 1.000 | 1.000 | 1.000 | 1.000 |
| 5 | 0.955 | 0.996 | 1.000 | 1.000 | 0.982 | 1.000 |
| 6 | 0.951 | 0.970 | 0.987 | 1.000 | 1.000 | 0.984 |
| 7 | 1.000 | 1.000 | 1.000 | 1.000 | 1.000 | 1.000 |
| 8 | 0.882 | 1.000 | 1.000 | 1.000 | 0.900 | 1.000 |
| 9 | 0.793 | 0.993 | 1.000 | 0.997 | 0.900 | 0.997 |
| 10 | 0.894 | 1.000 | 1.000 | 1.000 | 1.000 | 1.000 |
| 11 | 1.000 | 1.000 | 1.000 | 1.000 | 1.000 | 1.000 |

**Table 18:**
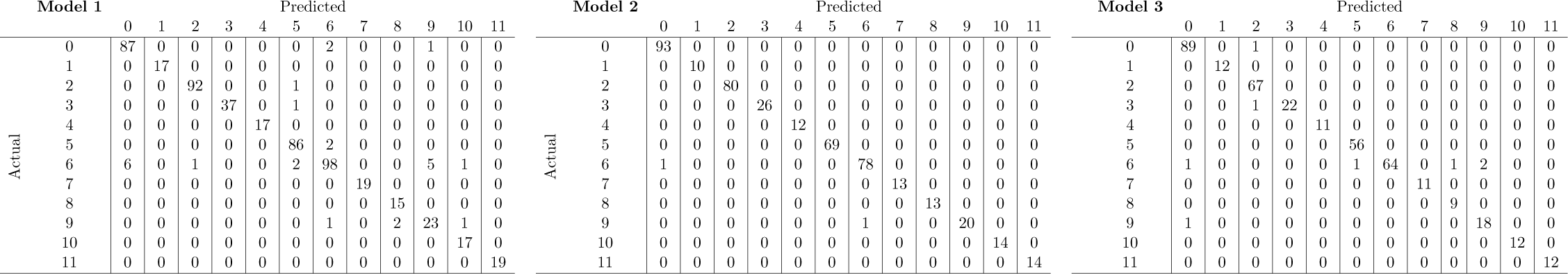
Confusion matrix for 12-class canine sarcoma classification.

### Public MS datasets

**Table 19:** Overall global accuracy, sensitivity, and specificity values for 2-class human ovary 1 classification.

| Model 2 |  | Transfer learning |  |
| --- | --- | --- | --- |
| Accuracy = $0.78 \pm 0.02$ | | Accuracy = $0.98 \pm 0.00$ | |
| Sensitivity | Specificity | Sensitivity | Specificity |
| 0.809 | 0.769 | 1.000 | 0.967 |

**Table 20:**
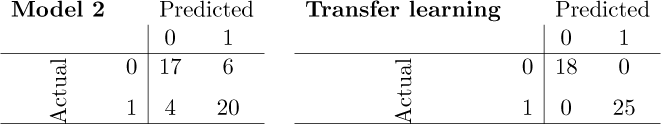
Confusion matrix for 2-class human ovary 1 classification.

**Table 21:** Overall global accuracy, sensitivity, and specificity values for 2-class human ovary 2 classification.

| Model 2 |  | Transfer learning |  | Cumulative representation learning |  |
| --- | --- | --- | --- | --- | --- |
| Accuracy = $0.803 \pm 0.009$ | | Accuracy = $0.833 \pm 0.021$ | | Accuracy = $0.996 \pm 0.001$ | |
| Sensitivity | Specificity | Sensitivity | Specificity | Sensitivity | Specificity |
| 0.933 | 0.750 | 1.000 | 0.735 | 1.000 | 1.000 |

**Table 22:** Confusion matrix for 2-class human ovary 2 classification.

| Model 2 |  | Predicted |  | Transfer learning |  | Predicted |  | Cumulative representation learning |  | Predicted |  |
| --- | --- | --- | --- | --- | --- | --- | --- | --- | --- | --- | --- |
|  |  | 0 | 1 |  |  | 0 | 1 |  |  | 0 | 1 |
| Actual | 0 | 14 | 9 | Actual | 0 | 20 | 9 | Actual | 0 | 20 | 0 |
|  | 1 | 1 | 27 |  | 1 | 0 | 25 |  | 1 | 0 | 29 |

### Comparison of our 1D-CNN against other ML approaches

**Table 23:** Optimized hyper-parameters for each ML algorithm.

| SVM | RF | LDA |
| --- | --- | --- |
| $C \in [0.1 - 0.8]$ | $ntree \in [50 - 1000]$ | $nu \in [2 - 10]$ |
| $\sigma \in [0.01 - 0.2]$ | $mtry \in [5 - 10]$ | $tol \in [0 - 10]$ |
| Kernel : radial basis function | node size $\in [10 - 50]$ | |

**Table 24:** Overall accuracy, sensitivity, and specificity values for 2-class canine sarcoma classification.

| SVM |  | RF |  | LDA |  |
| --- | --- | --- | --- | --- | --- |
| Accuracy = $0.76 \pm 0.16$ | | Accuracy = $0.96 \pm 0.01$ | | Accuracy = $0.88 \pm 0.17$ | |
| Sensitivity | Specificity | Sensitivity | Specificity | Sensitivity | Specificity |
| 0.465 | 1.000 | 0.850 | 0.994 | 0.639 | 0.947 |

**Table 25:**
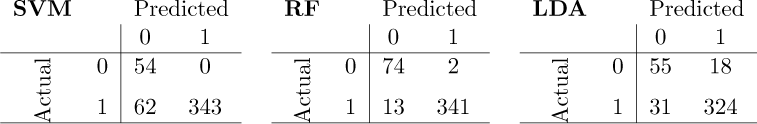
Confusion matrix for 2-class canine sarcoma classification.

**Table 26:**
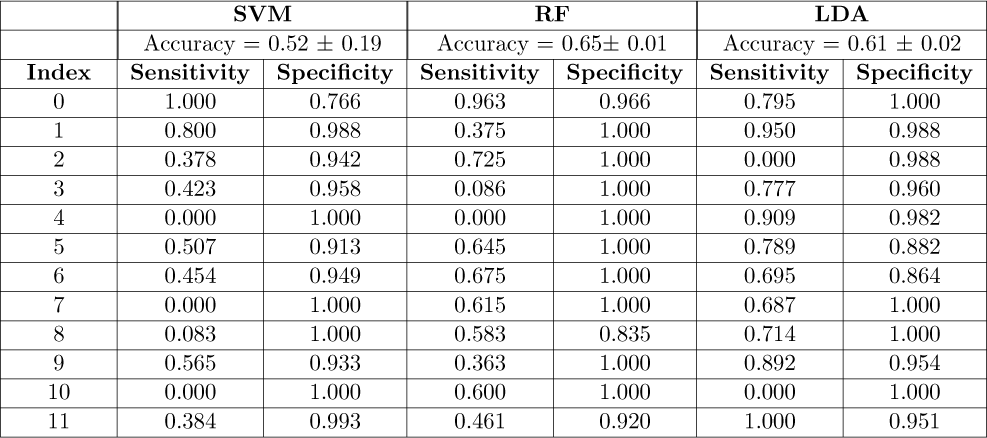
Overall accuracy, sensitivity, and specificity values for 12-class canine sarcoma classification.

**Table 27:**
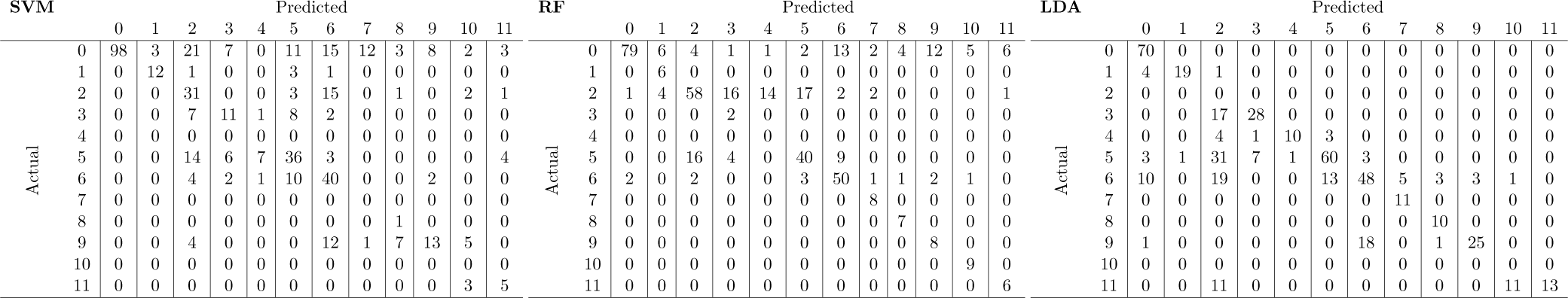
Confusion matrix for 12-class canine sarcoma classification.

**Table 28:** Overall accuracy, sensitivity, and specificity values for 3-class microorganisms classification.

|  | SVM |  | RF |  | LDA |  |
| --- | --- | --- | --- | --- | --- | --- |
| | Accuracy = $0.87 \pm 0.04$ | | Accuracy = $0.95 \pm 0.02$ | | Accuracy = $0.87 \pm 0.07$ | |
| Index | Sensitivity | Specificity | Sensitivity | Specificity | Sensitivity | Specificity |
| 0 | 0.964 | 0.807 | 0.922 | 0.914 | 0.777 | 1.000 |
| 1 | 0.720 | 0.975 | 0.853 | 0.955 | 1.000 | 0.916 |
| 2 | 0.930 | 1.000 | 1.000 | 0.993 | 1.000 | 0.928 |

**Table 29:**
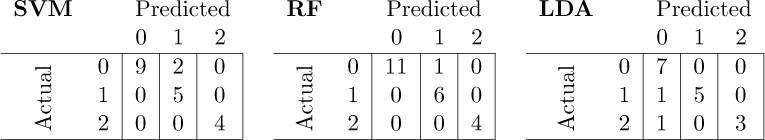
Confusion matrix for 3-class microorganisms classification.

**Table 30:** Overall accuracy, sensitivity, and specificity values for 5-class microorganisms classification.

|  | SVM |  | RF |  | LDA |  |
| --- | --- | --- | --- | --- | --- | --- |
| | Accuracy = $0.54 \pm 0.35$ | | Accuracy = $0.86 \pm 0.01$ | | Accuracy = $0.51 \pm 0.26$ | |
| Index | Sensitivity | Specificity | Sensitivity | Specificity | Sensitivity | Specificity |
| 0 | 1.000 | 0.588 | 1.000 | 0.933 | 0.625 | 1.000 |
| 1 | 0.000 | 1.000 | 1.000 | 1.000 | 0.000 | 1.000 |
| 2 | 0.667 | 1.000 | 0.666 | 0.937 | 0.750 | 0.615 |
| 3 | 0.000 | 1.000 | 0.666 | 1.000 | 1.000 | 1.000 |
| 4 | 1.000 | 0.842 | 1.000 | 1.000 | 0.400 | 0.800 |

**Table 31:**
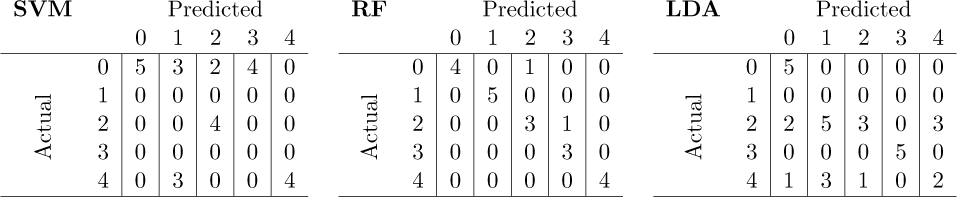
Confusion matrix for 5-class microorganisms classification.

**Table 32:**
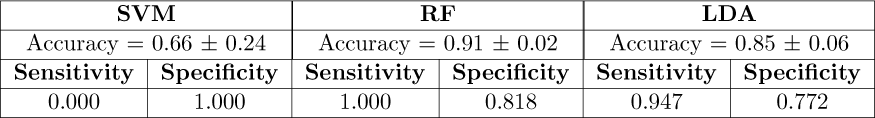
Overall accuracy, sensitivity, and specificity values for 2-class human ovary 1 classification.

| SVM |  | RF |  | LDA |  |
| --- | --- | --- | --- | --- | --- |
| Accuracy = $0.66 \pm 0.24$ | | Accuracy = $0.91 \pm 0.02$ | | Accuracy = $0.85 \pm 0.06$ | |
| Sensitivity | Specificity | Sensitivity | Specificity | Sensitivity | Specificity |
| 0.000 | 1.000 | 1.000 | 0.818 | 0.947 | 0.772 |

**Table 33:**
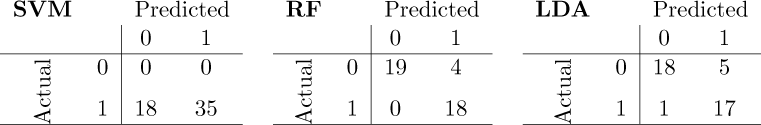
Confusion matrix for 2-class human ovary 1 classification.

**Table 34:** Overall accuracy, sensitivity, and specificity values for 2-class human ovary 2 classification.

| SVM |  | RF |  | LDA |  |
| --- | --- | --- | --- | --- | --- |
| Accuracy = $0.60 \pm 0.05$ | | Accuracy = $0.88 \pm 0.03$ | | Accuracy = $0.97 \pm 0.00$ | |
| Sensitivity | Specificity | Sensitivity | Specificity | Sensitivity | Specificity |
| 0.000 | 1.000 | 0.761 | 0.968 | 1.000 | 0.963 |

**Table 35:**
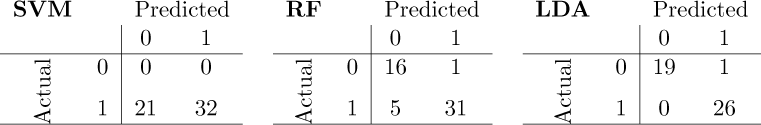
Confusion matrix for 2-class human ovary 2 classification.

